# A multipotent mouse distal airway secretory cell population that contributes to both airway and alveolar regeneration

**DOI:** 10.64898/2026.11.04.764405

**Authors:** Ullas V. Chembazhi, Shanru Li, Yun Ying, Zoey A. Miller, Chen Shen, Mijeong Kim, Subham Basu, Su Zhou, Michael P. Morley, Sarah E. Schaefer, Edward E. Morrisey

## Abstract

The distal airway is a spatially distinct compartment whose contribution to lung repair remains poorly defined. We have identified an epithelial cell population enriched in the most distal airways of the mouse lung, that is not present in the alveolar epithelium. Using lineage tracing tools based on Cldn10 expression which marks these cells, we show that the Cldn10+ lineage in homeostatic conditions marks distal airway secretory cells in a highly specific manner, with only a small contribution to multi-ciliated cells and no contribution towards the basal stem cell lineage. Remarkably, these Cldn10+ distal secretory cells proliferate and display a high degree of multi-potency across the lung epithelial axis after injury, generating basal cells, multi-ciliated cells, and alveolar epithelium. In severely remodeled regions after injury, we show that *Cldn10* lineage-labeled cells migrate from the airways and differentiate into alveolar epithelial cells. These findings establish the *Cldn10* lineage that defines a distal airway secretory facultative progenitor with both intrapulmonary airway and alveolar potential, providing a genetic system for studying distal lung repair.

## Introduction

The lung epithelium is largely quiescent under homeostatic conditions but mounts a substantial regenerative response following injury^1^. Respiratory viral infection, including influenza A virus, destroys both distal airway and alveolar epithelium, compromising barrier integrity and gas exchange^2–6^. Recovery requires that surviving epithelial cells proliferate, migrate into denuded regions, and differentiate to replace the lost cell types. When these programs are incomplete or misdirected, the result is persistent dysplastic remodeling rather than functional repair, a common feature of severe viral pneumonia and chronic lung diseases, including chronic obstructive pulmonary disease and idiopathic pulmonary fibrosis^7,8^. Identifying which cells carry out repair and what determines the fates they adopt is therefore central to understanding why regeneration succeeds in some settings and fails in others.

Epithelial progenitor activity is regionally organized along the proximal-distal axis of the lung^2,9–11^. Basal cells maintain the trachea and proximal airways, self-renewing and generating secretory and ciliated progeny^12,13^. Alveolar type 2 (AT2) cells act as the primary facultative progenitors in the alveolus, proliferating after injury and differentiating into alveolar type 1 (AT1) cells through transitional intermediates^14^. Between these compartments lies the bronchiolar epithelium, dominated by *Scgb1a1*^+^ secretory club cells, which function as facultative progenitors to repopulate the airway after ablation and have been reported to contribute to alveolar repair following influenza or bleomycin injury^15–19^. The magnitude and mechanisms of that alveolar contribution, however, remain unclear, in part due to a lack of specific mouse tools for studying this region^15,20^. Scgb1a1-driven alleles label secretory cells along the entire proximodistal axis and additionally mark a subset of AT2 cells at baseline, confounding the origins of alveolar lineage-labeled cells that arise from them during repair after injury^15,20^.

Compared to mice, humans have a distinct distal airway architecture in which terminal bronchioles transition through respiratory bronchioles before terminating into the alveolar space^21,22^. Respiratory bronchioles contain interwoven alveolar sacs and specialized Respiratory Airway Secretory (RAS) cells^22–24^. Unlike this, terminal bronchioles in mice open abruptly into alveolar ducts^21^. In mice, the distal bronchioles and bronchioalveolar duct junction (BADJ) mark the airway regions most proximate to the alveolar surface and are the most likely contributors of airway-derived alveolar repair^21^. Here we identify *Cldn10* as a secretory-restricted marker in the mouse lung epithelium and generated *Cldn10*^CreERT^^2^ to demonstrate that *Cldn10* lineage-labeled cells are concentrated in the distal airways, with no detectable overlap with AT2 cells at baseline. Following influenza infection, *Cldn10* lineage contributes to multilineage repair, forming contiguous patches of lineage-labeled cells in the distal airways, as well as alveolar cell types in the severely remodeled regions. Tracheal basal cells in organoids can give rise to Cldn10-expressing cells, and Cldn10 lineage-derived cells from mice in turn form 3D organoids that activate the *Sftpc* reporter under alveolar differentiation conditions. Together, these data define the distal airway secretory cell as an injury-responsive progenitor with both airway and alveolar competence and provide a genetic system for dissecting the signals that direct its fate.

## Results

### *Cldn10^CreERT2^* labels a secretory cell population enriched in the distal airways

To investigate secretory cell fates within the lung epithelial compartment during homeostasis, we isolated EpCAM-expressing epithelial cells from the lungs of adult CD1 mice and profiled them using single-cell RNA sequencing (scRNA-seq) to identify distinct gene expression programs **(Fig. 1A)**. Secretory cells robustly clustered and showed high expression of *Scgb1a1*. We identified that unlike *Scgb1a1,* which is expressed widely across distinct epithelial compartments **(Supplementary Fig. 1A)**, *Cldn10* is specifically expressed in the secretory cells **(Fig. 1A)**. We leveraged this finding and generated a new Cre recombinase-expressing mouse line driven by *Cldn10* (*Cldn10^CreERT2^*) **(Fig. 1B)**. We then crossed this Cre line to *ROSA26^tdTomato^ (tdTm)* to generate *Cldn10^CreERT2^; ROSA26^tdTomato^* to enable efficient lineage labeling and cell tracking. Induction with an optimized dose of tamoxifen for lineage labeling, followed by immunohistochemistry (IHC) seven days after induction, revealed that *Cldn10^+^* secretory cells in the adult mice were largely localized to the distal airways, with limited and sporadic presence in proximal airways **(Fig. 1C)**. This is strikingly distinct from the most widely used *Scgb1a1-CreER^TM^* mouse model for studying club cells, which labels cells across the entire proximal-distal axis of the airways.

**Figure 1.**
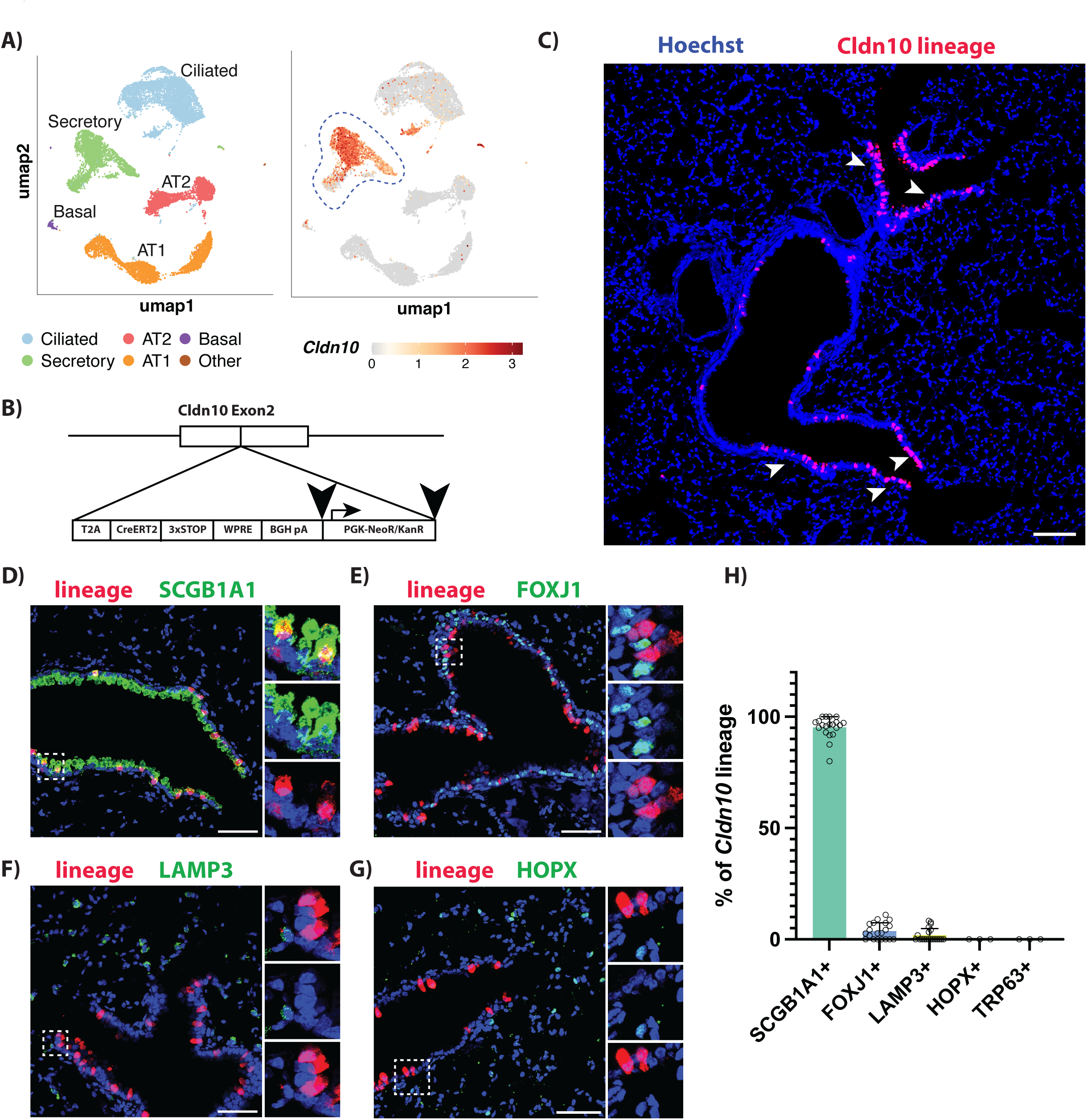
Cldn10-CreERT2 mice label distal secretory cells. **A)** scRNA-seq of EPCAM^+^ mouse epithelial cells from WT mice lung demonstrating that *Cldn10* is expressed specifically in the secretory cell compartment. **B)** CreERT2 was inserted into exon 2 of *Cldn10* gene to generate *Cldn10^CreERT2^* mice allele. **C-D)** *Cldn10^CreERT2^*labels distal airway bronchiolar region (C), specifically labeling secretory cells in this niche (D)**. E-G)** A minor fraction of ciliated cells is also labeled(E), with virtually no labeling of alveolar (F) AT2s or (G) AT1 lineages. **(H)** Quantitation showing labeling specificity of *Cldn10^CreERT2^*across different lung epithelial cell lineages. Unless stated otherwise, scale bars represent 50 um. Each dot represents fields of IHC images, taken across at least n=3 adult mice.

We next investigated the specificity of cell types labeled by *Cldn10^CreERT2^* by performing IHC along with markers of club cells (SCGB1A1), ciliated cells (FOXJ1), AT2 cells (LAMP3), AT1 cells (HOPX) and basal cells (TRP63) **(Fig. 1D-H)**. *Cldn10^CreERT2^* specifically labeled secretory cells in the airways (*Cldn10*^+^ cells) and did not co-stain with TRP63^+^ basal cells or HOPX^+^ AT1 cells **(Fig. 1F,G)**. We observed that a minor subpopulation of FOXJ1^+^ ciliated cells—which are largely known to be terminally differentiated^25^—were also labeled tdTm-positive upon tamoxifen induction in these mice **(Fig. 1E, H)**. We noticed a very small number of *Cldn10^CreERT2^*-labelled cells co-expressing *SFTPC* in the bronchioalveolar duct junction (BADJ), indicating labeling of bronchioalveolar stem cells (BASCs)^26^ **(Supplementary Fig. 1B)**. However, unlike the *Scgb1a1-CreER^TM^*mouse model, we did not observe co-labeling with SFTPC^+^/LAMP3^+^ AT2 cells in the alveolar region **(Supplementary Fig. 1A-C)**.

### Cldn10 lineage contributes to airway repair

To investigate the role of *Cldn10^+^* cells in distal airway repair after injury, we administered intranasal H1N1 PR8 influenza virus to *Cldn10^CreERT2^; ROSA26^tdTomato^* treated with tamoxifen 14-days prior to induction, which produced widespread injury in the mouse lung, comparable to human influenza infections **(Fig. 2A)**. We observed that 28 days post injury(dpi) that Cldn10 lineage-labelled cells increased in numbers, particularly in the most distal regions of the airway (**Fig. 2B,C**). Strikingly, a large fraction of these were seen as contiguous patches of lineage-labeled cells, that appear to re-establish lost airway epithelium (**Fig. 2B**).

**Figure 2.**
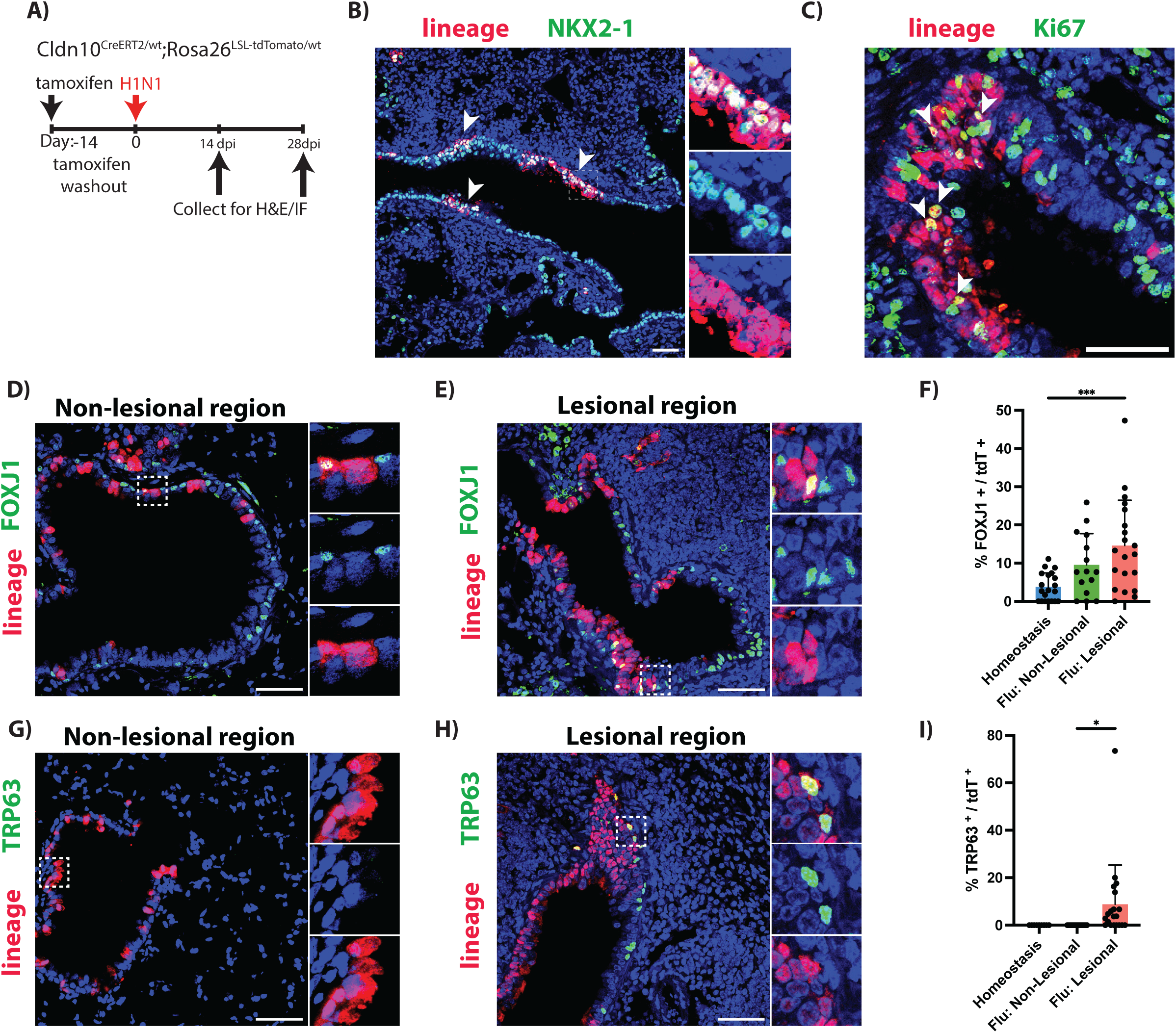
*Cldn10* lineage-derived cells contribute to distal airway repair. **A)** Schematic showing the schedule of influenza injury. **B-C)** *Cldn10* lineage cells proliferate in the distal airways, forming patches of lineage-derived cells at 14 dpi. Patches of Cldn10 lineage cells contain **(D-F)** FOXJ1^+^ ciliated and **(G-I)** TRP63^+^ basal-like identities at 28 dpi. These identities are prominent in severely remodeled (lesional) regions after injury (F and I). Unless stated otherwise, scale bars represent 50 um. Each dot represents fields of IHC images, taken across at least n=3 adult mice.

Next, we used the IHC approach to examine the identity of *Cldn10* lineage cells after severe influenza injury and to determine whether they contribute to different airway cell types (**Fig. 2D-I**). We noticed a considerable increase in the number of lineage-positive cells co-expressing FOXJ1, indicating differentiation toward a ciliated cell identity (**Fig. 2D-F**). Interestingly, we also noticed that a measurable population of *Cldn10* lineage cells was positive for the basal cell marker TRP63 (**Fig. 2G-I**).

### Cldn10 lineage contributes to alveolar lineages after influenza

To investigate if *Cldn10^+^* cells directly contribute to the re-establishment of alveolar epithelium after injury, we first evaluated if Cldn10-lineage cells migrate out of the airways. We found that at 28 days post influenza injury, *Cldn10* lineage cells delocalize from the airways and undergo significant migration into the severely damaged regions of the lung (**Fig. 3A,B**). In contrast, in the less severely injured regions, Cldn10 lineage cells remained largely attached to airways. Strikingly, we noticed that while airway-associated Cldn10 lineage cells largely expressed SCGB1A1 (**Fig. 3B**), maintaining secretory cell identity, the cells that get dislocated from the airways significantly downregulate SCGB1A1 expression (**Fig. 3B**). This implied that *Cldn10* secretory cells in the severely damaged regions migrate out of the airways into the alveoli, in severely damaged regions with simultaneous loss of secretory cell identity (**Fig. 3C**).

**Figure 3.**
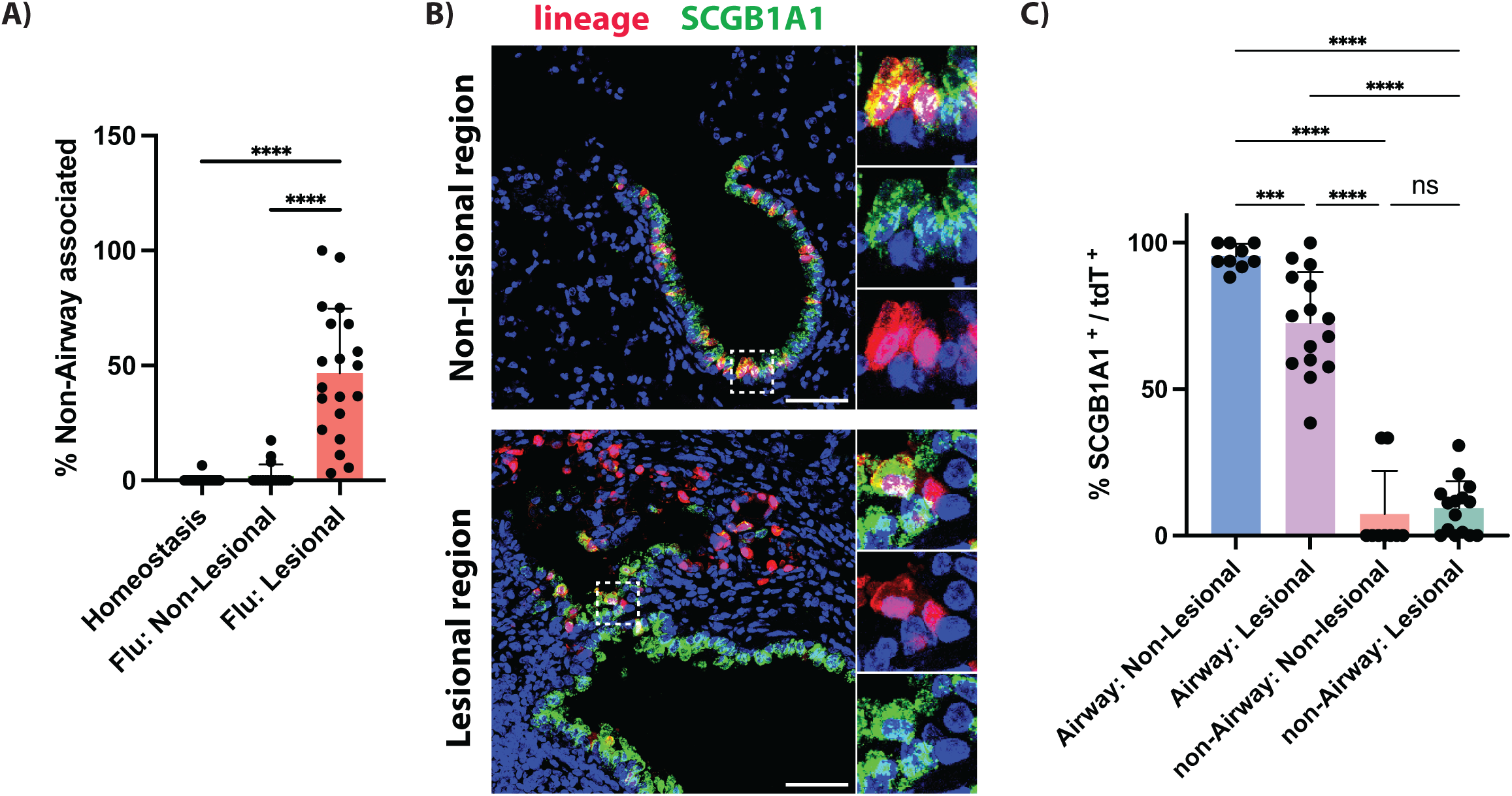
Airway-detached *Cldn10* lineage cells lose secretory cell identity. *Cldn10* lineage cells in severely remodeled (lesional) regions become **(A)** detached from airways, **(B-C)** downregulate SCGB1A1 expression, and localize to the non-airway regions. Unless stated otherwise, scale bars represent 50 um. Each dot represents fields of IHC images, taken across at least n=3 adult mice.

We next investigated whether Cldn10 lineage cells acquire alveolar identities after injury. As expected, Cldn10 lineage cells in the less severe regions remained largely SCGB1A1^+^, airway-associated and did not express markers of AT1 or AT2 cells (**Fig. 4A,D**). In contrast, in the severely remodeled regions, we observed a significant increase in the number of *Cldn10* lineage-derived LAMP3-expressing cells (**Fig. 4B,C**). We noticed that airway-associated Cldn10 cells largely retained airway identities in severely injured regions; the SFTPC-expressing lineage-positive population largely comes from cells that are no longer associated with the airways (**Fig. 4B,C**). To assess whether the alveolar upregulated populations differentiate into AT1 cells, we stained the cells with the AT1 marker *HOPX* (**Fig. 4D-F**). We noticed that a small fraction of *Cldn10* lineage-positive cells upregulates HOPX expression (**Fig. 4E,F**), suggesting they may gain an AT1-like identity. Whether *Cldn10* lineage-derived cells transition into fully functional alveolar epithelial cells and contribute to gas exchange remains to be determined.

**Figure 4.**
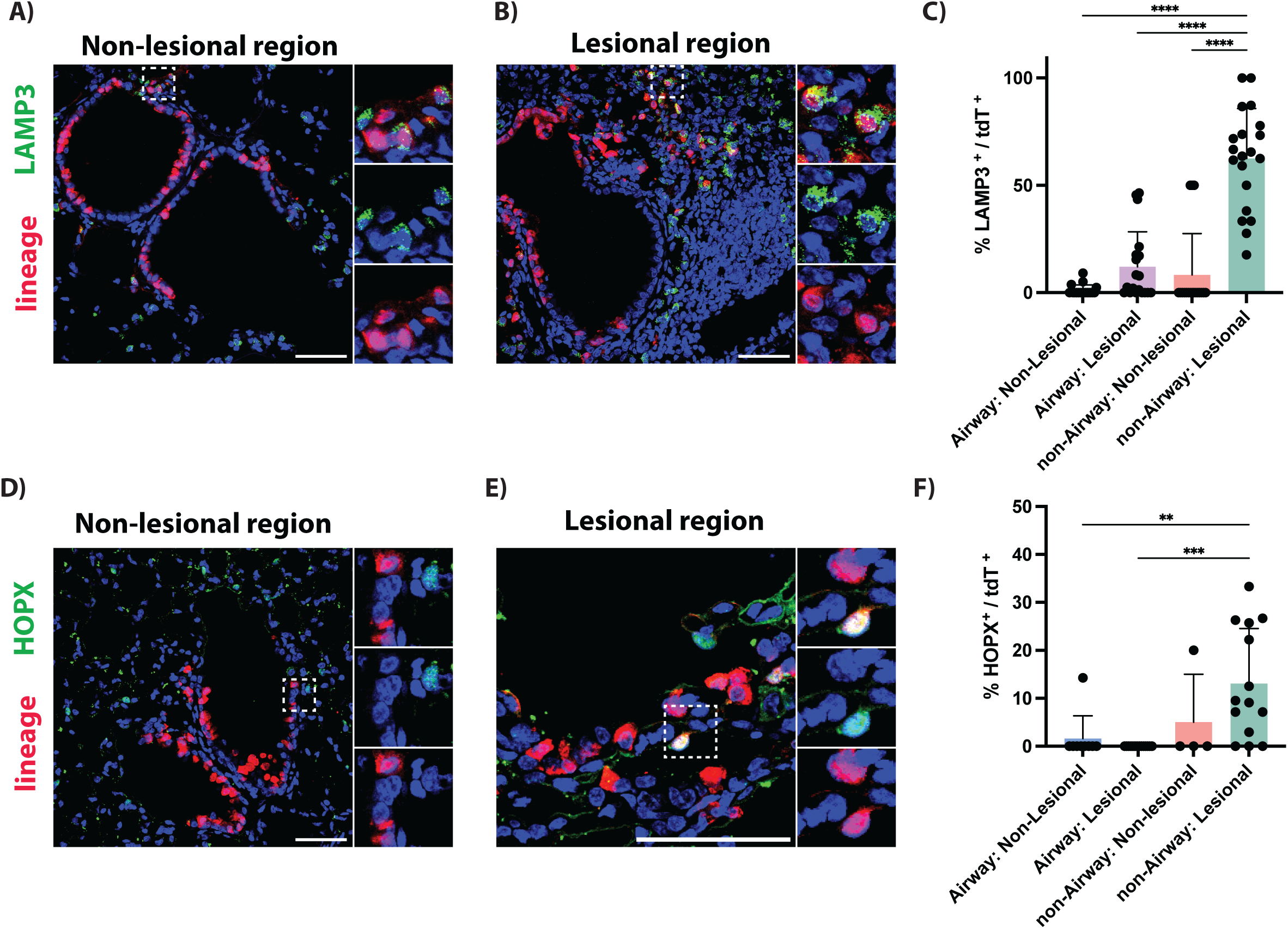
Airway-detached *Cldn10* lineage gain alveolar cell identities. **A)** Lineage-derived cells remain airway-attached in non-lesional regions without a prominent increase in LAMP3 expression, in contrast to **(B-C)** lesional regions where a significant number of lineage-derived cells are found detached from the airways with upregulated AT2 marker expression. **D-F)** A similar trend is seen with expression of HOPX, potentially suggesting AT1-like identity. Unless stated otherwise, scale bars represent 50 um. Each dot represents fields of IHC images, taken across at least n=3 adult mice.

### Cldn10+ progenitors can be derived from basal cell organoids

To investigate the progenitor potential of *Cldn10*-traced cells, we isolated *EpCAM*^+^*tdTm^+^*cells from adult *Cldn10^CreERT2^; ROSA26^tdTomato^* mice one week after induction with tamoxifen and plated them in Matrigel. We observed that under airway media conditions (see methods), *Cldn10* lineage-derived cells formed 3D organoids within three weeks in culture, indicating their progenitor potential (**Fig. 5A-C**). To investigate whether these cells can differentiate into AT2 cells, we leveraged *Cldn10^CreERT2^;Sftp^cEGFP^;ROSA26^tdTomato^* mice which enables live tracking of transition into AT2 fate *ex vivo* by expression of *Sftpc*-*IRES-EGFP* reporter. We isolated *EpCAM*^+^*tdTm^+^EGFP ^-^* mice one week after induction with tamoxifen and plated them in Matrigel under airway media conditions for two weeks and then transitioned them into alveolar media conditions (see methods) for another week. We observed robust EGFP expression, indicating Sftpc expression and a transition to an AT2-like fate (**Fig. 5C**).

**Figure 5.**
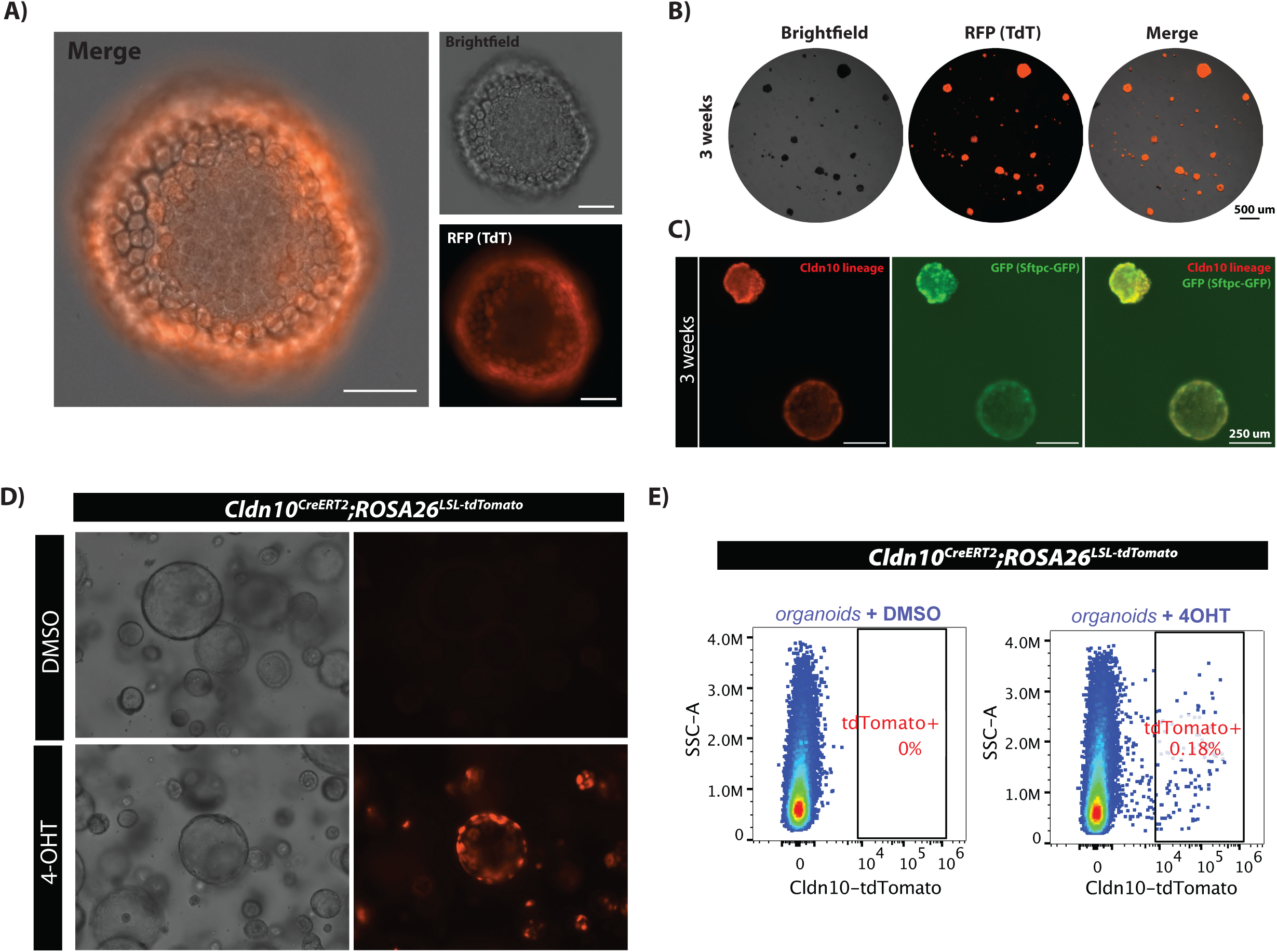
Cldn10 progenitors produce organoids that can differentiate into AT2s. A-B) *Cldn10* lineage derived cells produced 3D organoids in 2-3 weeks (**C)** and upregulate expression of an SFTPC-IRES-GFP upon transition to alveolar media conditions. **D-E)** Basal cells differentiate into *Cldn10* expressing cells in 3D organoid cultures. Unless stated otherwise, scale bars represent 50 um.

Basal cell organoids are known to differentiate into secretory and ciliated cells within 3D organoid cultures. As discussed, we observed that *Cldn10* lineage produced *TRP63^+^* basal cells after influenza injury (**Fig. 2H**). We next evaluated whether *Cldn10*-expressing cells can be derived from basal cells in culture using *Cldn10^CreERT2^* mice. To test this, we cultured *NGFR^+^EpCAM^+^* basal cells isolated from the trachea of uninduced adult *Cldn10^CreERT2^; ROSA26^tdTomato^* mice for 12 days and then were treated with 4-Hydroxytamoxifen (4-OHT) to activate CreERT2-driven recombination of *ROSA26^LSL-tdTomato^* locus in *Cldn10*-expressing cells, thereby labeling them specifically. We observed that a small fraction of tdTomato^+^ cells within basal cell organoids using live microscopy and flow cytometry, suggesting that basal cells can differentiate into *Cldn10*-expressing cells in *ex vivo* organoid cultures (**Fig. 5D, E**).

## Discussion

Previous studies have shown that airway cell types can generate alveolar epithelial cells upon acute injury. These studies are based on either the Scgb1a1 or Sox2 inducible lineage-tracing systems, which are not specific to any particular lineage. In particular, the *Scgb1a1-CreER^TM^* line marks an appreciable number of AT2 cells and labels various other airway lineages, confounding the ability to track airway contribution to other cell types using this gene as a lineage tracing driver^15,20,27^. Here we describe a new mouse lineage fate mapping allele, *Cldn10^CreERT2^*, which addresses these significant limitations and reveals a spatial specificity restricted to the distal airway. Using this new model, we show that Cldn10+ distal airway epithelium is primarily composed of secretory cells with a small number of multi-ciliated cells in the distal airway, along with rare BASCs. Upon injury, Cldn10+ cells generate a full spectrum of airway epithelium across the proximal-distal axis, including basal cells and AT2 cells. These studies reveal the high level of plasticity across the epithelial compartment in the lung in a rigorous manner, providing a new model to examine the lineage barriers between the various epithelial lineages in maintaining lung health and disease prevention.

We note that Cldn10 lineage-labeled cells that appear to detach from the airway in severely remodeled regions downregulate SCGB1A1 and acquire LAMP3 and, in a smaller fraction, HOPX expression. Within the same lungs, labeled cells in less severely injured territory remain airway-associated and retain secretory identity, implying that transition to alveolar conversion is driven by the local injury environment. This is consistent with facultative behavior described for other lung epithelial progenitors^28–31^. While lineage-labeled cells gain LAMP3/HOPX expression, it remains to be determined if they become fully differentiated alveolar cell types and form functional alveolar units that participate in gas exchange, or if they remain a bipotent AT1/AT2 intermediate^30,32–34^.

We show that *Cldn10* lineage generates TRP63+ cells within severely injured airways. Basal cells are largely restricted to the trachea and proximal airways in mice under homeostatic conditions^13,35^, and their appearance in the alveolar region after influenza injury has been associated with dysplastic remodeling rather than productive repair^36^. The lineage-labeled basal cells we observe occur within contiguous patches that also contain secretory and ciliated cells, which is more consistent with reconstitution of a stratified airway epithelium than with the dysplastic pods described previously, but it remains to be assessed whether they resolve or persist. Future research will address whether distal secretory cells adopting a basal program represent a repair intermediate or an early step toward dysplastic remodeling.

Our finding that tracheal basal cells cultured gave rise to *Cldn10^+^* cells indicates that this population can be generated from basal cells. Alongside our observation that the *Cldn10* lineage can give rise to TRP63^+^ progeny *in vivo*, this suggests a reciprocal relationship with the basal compartment, where secretory and basal identities in the airway are interconvertible under defined conditions, consistent with the plasticity described in other airway injury models^2,16,37^. Further, *Cldn10* lineage-derived cells formed 3D organoids, and activated expression of the *Sftpc* reporter under alveolar differentiation conditions, highlighting their progenitor potential and indicating that their alveolar competence is cell-intrinsic and can be elicited *ex vivo*.

It must be noted that the regional distribution we report is based on histological assessment of lineage-labeled cells, reflecting both endogenous expression patterns and recombination efficiency. We also observed labeling of a minor FOXJ1^+^ population at baseline, representing *Cldn10* expression in ciliated cells. However, we expect minimal confounding effects from this, as ciliated cells are generally considered terminally differentiated. In this study, we focused on influenza infection as a physiologically relevant model of severe distal lung injury. Future studies will be needed to determine whether the cellular behaviors described here are shared across other injury models, such as bleomycin-induced damage, and to define how this repair process evolves over time.

Together, these findings identify the *Cldn10* lineage distal airway secretory cell as an injury-responsive progenitor with competence for both airway and alveolar fates. *Cldn10^CreERT2^* provides a highly specific platform to study repair processes within the airway and their contribution to the alveolar compartment, given the absence of AT2 labeling. Distal-enriched labeling of *Cldn10^CreERT2^* permits interrogation of the distal airway as a distinct repair niche, which is of particular interest given that the human distal airways contain secretory populations without known mouse counterparts, that are transcriptionally distinct from proximal airways. The mouse model described here now provides a way to identify the niche-specific mechanisms governing distal airway secretory cell fate — a necessary step toward promoting functional regeneration.

## Methods

### Animals

All mouse experiments were performed following the protocols of the University of Pennsylvania Institutional Animal Care and Use Committee. 2– 3-month-old adult mice were used in all the experiments. Mice were housed in groups of up to five animals per cage. The following mice were obtained from the Jackson Laboratories: *ROSA26^LSL-tdTomato^* (Gt(ROSA)26Sortm14(CAG-tdTomato)Hze, strain #007914)^38^ and *Scgb1a1-CreER^TM^* (B6N.129S6(Cg)- Scgb1a1tm1(cre/ERT)Blh/J)^15^. Mice were maintained on a mixed genetic background.

*Cldn10^CreERT2^* mice was created in this study. Using CRISPR-based homology-directed repair in mouse embryonic stem cells, a tamoxifen-inducible *CreERT2* sequence was inserted in the exonic region of Cldn10 coding sequence after a T2A cleavage sequence. This sequence is followed by a 3xSTOP sequence. An FRT-flanked Neomycin resistance cassette was used to select positive clones during ES cell targeting. Flp-mediated recombination using the *ROSA26^FlpO^* mouse allele (JAX strain #012930) was used to remove the FRT-flanked Neo cassette. This allows expression of *CreERT2* under the control of the Cldn10 promoter.

### Tamoxifen administration

Tamoxifen (Sigma-Aldrich #T5648) was suspended in a 1:10 mixture of ethanol and corn oil to prepare a stock concentration of 20 mg/mL. Tamoxifen solution was delivered to mice by oral gavage of 10 μL per gram body weight for two or three consecutive days.

### Organoid culture

Single-cell suspension isolated by FACS was pelleted, resuspended in Matrigel (Corning), and plated as domes (targeting 500 cells/ul whenever possible). EPCAM+NGFR+ basal cells and EPCAM+tdTomato+ Cldn10-lineage cells were cultured in airway media based on complete serum-free differentiation medium (cSFDM)^39^. For organoid differentiation experiments, airway media were replaced with alveolar media conditions as described previously^40^.

### Influenza injury

For Influenza injury, mice were infected with A/Puerto Rico/8/34 H1N1 influenza virus (ATCC #VR-95PQ, Lot 70022722) at a dose determined empirically in our laboratory to cause 15-20% weight loss without death (1:6,000,000) as previously reported^41^. In all cases, H1N1 was diluted in sterile saline and administered intranasally in a volume of 50 μL. This resulted in a moderate level of tissue damage with heterogeneous zones of injury. Mice were weighed daily for two weeks following infection and euthanized if their weight loss exceeded 30% of their original body weight.

### Histology

For tissue preparation, mice were euthanized by a lethal dose of CO_2_ inhalation, and the lungs were perfused with cold PBS, followed by inflation with cold 2% Paraformaldehyde at a constant pressure of 25 cm H_2_O. Tissues were fixed overnight at 4°C in 2% Paraformaldehyde and dehydrated in a series of ethanol concentration gradients. Tissues were then embedded in paraffin and sectioned at 6 μm thickness. Hematoxylin and eosin (H&E) and Immunohistochemistry (IHC) assays were performed as previously described^8,16^. Fluorescence images were acquired at 20x and 40x using z-stacks on a Zeiss LSM 710 or a Leica Stellaris 5 laser-scanning confocal microscope. Cell counting analysis was performed using Qpath and FIJI.

### Single-cell isolation, flow cytometry, and FACS

Lungs were harvested and digested into single-cell suspensions as previously described^8^. Briefly, lungs were perfused with cold PBS through the right ventricle, minced with a razor blade, and digested in a buffer supplemented with collagenase I, dispase, and DNase I. After trituration and filtration, cells were stained with a combination of antibodies on ice for 30 min for flow cytometry and FACS. To enrich epithelial retrieval, CD45+ and CD31+ cell populations were depleted from the cell suspension using MACS kits (Miltenyi Biotec) according to the manufacturer’s protocol.

### Mouse single-cell RNA sequencing

scRNA-seq was performed using EPCAM^+^CD45^-^CD31^-^ single cells isolated from wild-type mice using 10X Genomics V4 chemistry. Cells were loaded onto a 10X Chromium Controller (10X Genomics), cDNA and sequencing libraries were prepared according to the manufacturer’s instructions, and cDNA/library quality control was performed using an Agilent 2100 Bioanalyzer with the Bioanalyzer High Sensitivity DNA kit (Agilent #5067-4626). Sequencing was performed by Azenta Life Sciences on an Illumina NovaSeq X Plus system. Analysis was performed as previously described^41^. Briefly, reads were aligned to the mouse reference genome, unique molecular identifier (UMI) counts were obtained, ambient RNA contamination was removed, doublet detection was performed, and analysis was performed using Seurat in R^42^.

### Statistical analysis

In all mouse experiments, statistical significance was determined using one-way ANOVA in GraphPad Prism 11. Statistical analysis of scRNA-seq data was performed in R.

## Acknowledgements

We acknowledge and thank the core facilities at the University of Pennsylvania and Children’s Hospital of Philadelphia (CHOP) without whom we could not have completed this work. Flow cytometry data in this manuscript were generated at the CHOP Flow Cytometry Core Laboratory. Confocal microscopy was performed in the Penn Cell and Developmental Biology Microscopy Core. We thank the members of the Morrisey lab, Basil lab, and Penn-CHOP Lung Biology Institute (LBI) for their helpful suggestions and discussions throughout this work.

---

**Supplementary Figure 1.**
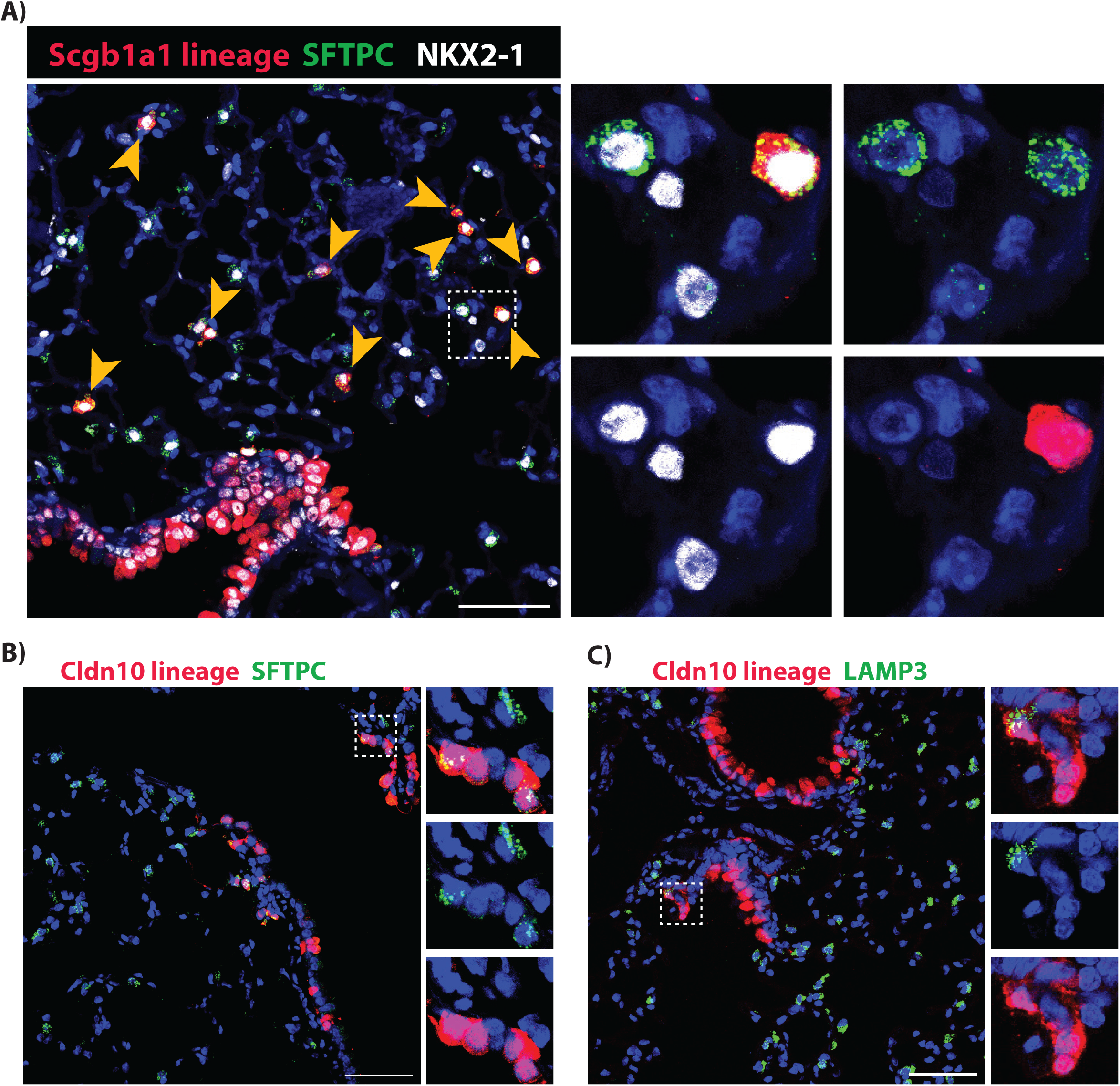
Cldn10CreERT2 mice label distal secretory cells. **A)** *Scgb1a1-CreER^TM^* labels several cell types, including alveolar cell types AT2s in homeostasis, confounding airway secretory cell studies. **B-C)** *Cldn10^CreERT2^*shows virtually no alveolar labeling, but labels cells at the BADJ, likely representing BASCs. Unless stated otherwise, scale bars represent 50 um.

